# Conscious perception and the modulatory role of dopamine: no effect of the dopamine D2 agonist cabergoline on visual masking, the attentional blink, and probabilistic discrimination

**DOI:** 10.1101/2020.03.17.994863

**Authors:** E.A. Boonstra, M.R. van Schouwenburg, A.K. Seth, M. Bauer, J.B. Zantvoord, E.M. Kemper, C.S. Lansink, H.A. Slagter

## Abstract

**Rationale:** Conscious perception is thought to depend on global amplification of sensory input. In recent years, striatal dopamine has been proposed to be involved in gating information and conscious access, due to its modulatory influence on thalamocortical connectivity.

**Objectives:** Since much of the evidence that implicates striatal dopamine is correlational, we conducted a double-blind crossover pharmacological study in which we administered cabergoline – a dopamine D2 agonist – and placebo to 30 healthy participants. Under both conditions, we subjected participants to several well-established experimental conscious-perception paradigms, such as backward masking and the attentional blink task.

**Results:** We found no evidence in support of an effect of cabergoline on conscious perception: key behavioral and event-related potential (ERP) findings associated with each of these tasks were unaffected by cabergoline.

**Conclusions:** Our results cast doubt on a causal role for dopamine in visual perception. It remains an open possibility that dopamine has causal effects in other tasks, perhaps where perceptual uncertainty is more prominent.

## Introduction

The relationship between consciousness and the brain is often lauded as one of the big mysteries in contemporary science. How does the brain constrain its own spontaneous activity as well as the influences it undergoes from outside, in the determination of conscious awareness? Several influential theories propose that consciousness is related to the ‘broadcasting’ of sensory information to the whole brain and that thalamocortical circuits serve as an important mediator of such broadcasting (Crick & Koch, 2003; Dehaene & Changeux, 2011; Edelman, 2003). The broadcasting of sensory information necessitates the occurrence of selection or filtering, simply because not everything which takes place in the brain reaches conscious awareness.

With the requirement of selection, we cannot ignore the role of the basal ganglia: a cluster of subcortical nuclei located deep in the brain, which modulate activity of thalamocortical circuits (Smith, Raju, Pare, & Sidibe, 2004) and have long been implicated in action selection; the decision to execute one of several possible behaviors (Redgrave, Prescott, & Gurney, 1999). Notably, the basal ganglia are connected through parallel loops via the thalamus not only to the motor cortex, but to many parts of frontal cortex as well (Alexander, DeLong, & Strick, 1986). Hence, it is capable of modulating a wide range of cognitive operations. Indeed, the basal ganglia have been implicated in working memory updating (Frank & O’Reilly, 2006), attention shifting (Cools, 2011), and visual categorization (Seger, 2008): cognitive acts that support suggestions concerning the common principle underlying the basal ganglia’s operations; namely, selection (Frank, Loughry, & O’Reilly, 2001; Redgrave, Prescott, & Gurney, 1999).

The striatum – the biggest structure constituting the basal ganglia – deserves special attention on this topic. As the basal ganglia’s primary input nucleus, the striatum is well-positioned to play a pivotal role in the basal ganglia’s selective functionalities, as terminal fields from different cortical regions converge in the striatum (Yeterian & van Hoesen, 1978; Haber, Kim, Mailly, & Calzavara, 2006; Mailly, Aliane, Groenewegen, Haber, & Deniau, 2013; Heilbronner, Meyer, Choi, & Haber, 2018). While dopaminergic projections are usually associated with reward prediction error (Schultz, 2016), and the role of reward in perceptual decision making (Ding & Gold, 2013); less well-studied signaling of the (dopamine-infused) striatum include saliency, threat, processing of sensory information, and promoting of behaviors reliant on sensory information (Cox & Witten, 2019).

Despite the suggestion that the basal ganglia may be “mute” as regards to consciousness (Boly et al., 2017), it has been known for a long time that basal ganglia structures are involved in sensory and perceptual processes (Alexander & Crutcher, 1990; Arsalidou, Duerden, & Taylor, 2013; L. L. Brown, Schneider, & Lidsky, 1997; Seger, 2013). Both the striatum and dopaminergic firing have been implicated in the tight relationship between perception and action, and consciousness more generally. For example, dopamine-depleted mice are unable to attend to salient sensory information and choose appropriate actions, suggestive of a critical role for dopamine in the expression of consciousness (Palmiter, 2011). In humans as well, it has recently been shown that minimally conscious patients suffer from a dopaminergic deficit in presynaptic neurons projecting to the striatum and central thalamus (Fridman, Osborne, Mozley, Victor, & Schiff, 2019).

In humans, the striatum and its irrigation by dopamine have also been implicated in well-known experimental paradigms used to study the neural correlates of consciousness, such as the attentional blink and backward masking task. In the attentional blink task, a deficit occurs when people have to detect two target stimuli (T1 and T2) presented in close temporal succession among distracter events. Specifically, when T2 follows T1 within 100– 500 ms, it often goes unnoticed. This deficit is called the attentional blink (AB; Shapiro, Raymond, & Arnell, 1997). Healthy participants with more D2-like receptor binding in the striatum – as shown with PET – showcased a larger AB (Slagter et al., 2012). In addition, intracranial EEG recordings in the ventral striatum revealed a short-latency increase in theta-band oscillatory activity only for consciously perceived target stimuli (Slagter et al., 2017).

In backward masking tasks, processing of target stimuli is interrupted by presenting a mask in close succession to the target (Breitmeyer, 2007). Studies employing fMRI consistently show differences in BOLD activity in the striatum and thalamus between seen and unseen stimuli using backward masking tasks (Bisenius, Trapp, Neumann, & Schroeter, 2015). In another PET study, dopamine D2 binding potential in the right striatum was found to correlate positively with both objective (task performance) and subjective (seen/unseen) visibility during backward masking (Van Opstal et al., 2014). These studies collectively suggest a role for the striatum and dopaminergic activity in the selection of visual information and the formation of conscious visual percepts.

However, up until now the relationship between striatal dopamine and conscious perception is based on correlational evidence. As such, in the present study we sought to manipulate this relationship experimentally, by administering the dopamine D2 agonist cabergoline to healthy participants. Out of the two main dopamine receptor families, D2 receptors have been found to be more prevalent in the striatum, while D1 receptors are present more in the prefrontal cortex (Gerfen, 1992). We chose cabergoline because it has greater affinity for D2 receptors, and it has been reported to have less side-effects compared to other D2 agonists such as bromocriptine (Frank & O’Reilly, 2006). Cabergoline, at low doses, has been suggested to preferentially stimulate pre-synaptic D2 autoreceptors, which have been found to inhibit phasic dopamine bursts in the striatum (Ford, 2014; Frank & O’Reilly, 2006). Cabergoline has been successfully administered in small dosages (1-1.5 mg) to manipulate performance in healthy participants on working memory tasks (Broadway, Frank, & Cavanagh, 2018; Fallon, Zokaei, Norbury, Manohar, & Husain, 2017; Frank & O’Reilly, 2006), a modified version of the Simon task (Cavanagh, Masters, Bath, & Frank, 2014), as well as action cancellation and error awareness tasks (Nandam et al., 2013). When cabergoline was administered to Parkinson’s patients for longer periods of time, decreases contrast sensitivity was found (Hutton, Morris, & Elias, 1999). We administered cabergoline in an attempt to manipulate performance on two paradigms traditionally used to study the neural correlates of conscious perception: backward masking and the attentional blink task. In addition, there is increasing evidence that dopamine plays a crucial role in determining the influence of sensory information in relation to expectations acquired through past experience (Cassidy et al., 2018; Friston et al., 2012). To investigate this relationship, we subjected participants to a probabilistic discrimination task in which the probability of stimulus occurrence varied across blocks of trials, thereby tapping into the learning capabilities the basal ganglia is implicated in traditionally (Berke, 2018).

The difficulty with manipulating dopamine is that effects have been found to depend on baseline dopamine levels in accordance with an inverted-U-shape (Cools & D’Esposito, 2011). This means that dopamine is thought to have an optimal level for task performance, but that dopamine manipulations may either benefit or worsen performance dependent on an individual’s starting point on the U-curve. Two measures used to estimate baseline dopamine levels are working memory operation span (OSPAN; Broadway et al., 2018; Cools, Gibbs, Miyakawa, Jagust, & D’Esposito, 2008) and spontaneous eye-blink rate (sEBR; Cavanagh et al., 2014; Jongkees & Colzato, 2016). We used these measures to analyze and control for potentially different effects of cabergoline in relation to baseline dopamine levels.

## Methods

### Participants

30 native Dutch-speakers were recruited from the University of Amsterdam subject pool to complete the experiment (mean age = 22, range 18-29, 25 female). Because our study is the first to investigate the effects of cabergoline on conscious perception, we did not conduct a power analysis. Instead, our sample size is on the upper end of sample sizes employed by previous research in which cognitive and neural effects of cabergoline were reported with 12-30 participants (Cavanagh et al., 2014; Cohen, Krohn-Grimberghe, Elger, & Weber, 2007; Frank & O’Reilly, 2006; Nandam et al., 2013; Norbury, Manohar, Rogers, & Husain, 2013; Yousif et al., 2016; Fallon et al., 2017; Broadway et al., 2018), and in which the behavioral and event-related potential (ERP) effects were reported which we aimed to manipulate with cabergoline (12-15 subjects; Del Cul et al., 2007; van Opstal et al., 2014; Slagter et al., 2012).

Four participants experienced adverse reactions during the cabergoline session (dizziness and nausea) that interfered with their ability to participate in the study. In two participants, nausea was present to the point of vomiting. One participant dropped out due to headaches in the placebo session. Completers and non-completers did not differ in baseline age, BMI, heart rate/blood pressure, or baseline dopamine proxies (see below). It should be noted however that all drop-outs were women.

In total, 124 individuals were considered for inclusion in the study. 113 were interviewed over the telephone to ensure normal or corrected-to-normal vision, no history of neurological, psychiatric, or any other relevant medical problems, and abstinence from psychoactive medication. The use of hormonal contraceptives served as an additional inclusion criterium for female participants, who were tested outside of their period due to variability in D2R availability during different phases of the menstrual cycle (Czoty et al., 2009). 63 individuals were eligible for participation, out of which 35 participants were available to schedule the required three lab visits (see Procedure). 5 participants did not meet requirements for inclusion based on the first lab visit. After excluding 5 participants with adverse reactions, the final sample consisted of 25 participants (mean age = 22, range 18-29, 20 female). In exchange for their participation, participants received 10 euros an hour, with a minimum of 110 euros in total. The study protocol was approved by the medical ethical committee of the Academic Medical Centre, Amsterdam (currently Amsterdam University Medical Centers). All participants provided written informed consent in accordance with the Declaration of Helsinki.

### Procedure

Participants came to the lab three times on different days (see Table 1). Once for screening (duration: 2.5h), and twice for an experimental session in which either placebo or 1.5mg of cabergoline was administered orally (duration: 4.5h each), as part of a double-blind crossover design. Previous studies found cognitive and neural effects of cabergoline using a dosage of 1-1.5mg (Cavanagh et al., 2014; Cohen et al., 2007; Frank & O’Reilly, 2006; Nandam et al., 2013; Norbury et al., 2013; Yousif et al., 2016; Fallon et al., 2017; Broadway et al., 2018). As with our sample size, we chose a dosage of 1.5mg to be on the upper end of previously employed dosages. There was at least a day in between screening and the first session, and at least a week between both sessions.

**Table 1.** Experimental procedures for the screening, placebo, and cabergoline sessions.

| Screening | Sessions (placebo/cabergoline) |
| --- | --- |
| Medical questionnaire | Questions regarding recent substance use + pregnancy test |
| M.I.N.I. | Blood pressure/heart rate + VAS 1 |
| Body weight/height | Drug intake |
| Blood pressure/heart rate | 40-minute break |
| sEBR | EEG setup |
| OSPAN | sEBR |
| Titration backward masking | Blood pressure/heart rate + VAS 2 |
| Titration probabilistic discrimination | Backward masking (± 1.5h after drug intake) |
|  | 30-minute lunch |
|  | Attentional blink |
|  | Reaction time task |
|  | Probabilistic discrimination |
| | Blood pressure/heart rate + VAS 3 (± 3.5h after drug intake) |

#### Screening

The first lab visit took place anywhere between 09:00 and 17:30. After providing written informed consent, participants answered a series of questions concerning potential medical conditions. Next, we conducted the M.I.N.I.; a structured screening interview for DSM-IV axis-I disorders (Sheehan et al., 1998). We subsequently measured participant’s weight, height, BMI, blood pressure (BP), and heart rate. Participants were included in the experiment only if these measures fell within pre-established bounds (BMI 18-30, diastolic BP < 50 or > 90 mmHg, systolic BP < 95 or > 140 mmHg). Next, six external electrodes were attached to the participant’s face and ears in order to measure spontaneous eye-blink rate (sEBR) at rest. Finally, participants completed an operation span (OSPAN) working-memory task (Unsworth, Heitz, Schrock, & Engle, 2005), and a titration procedure for two behavioral tasks to be completed during both experimental sessions (backward masking and probabilistic discrimination; see below).

#### Session

All placebo and cabergoline sessions (the second and third visit) took place between 08:30 and 14:00. Participants were instructed to abstain from drug and heavy alcohol use, the day before and during the day of the session. Also, participants were instructed to abstain from caffeine and nicotine the morning of the session. Compliance to the instructions was checked by the examiner on arrival, in case of non-compliance the session was postponed. Female participants completed a midstream pregnancy test. Breakfast was offered, in order to avoid cabergoline-intake on an empty stomach. Blood pressure and heart-rate were measured using an Omron® M3 comfort Sphygmomanometer, and participants filled in a visual analogue scale (VAS, see below) three times during the session: on arrival, at around 1.5h after placebo or cabergoline intake, and at the end of the session. After the initial blood pressure/heart-rate/VAS measurement, participants were administered either placebo or cabergoline in a double-blind fashion (order randomized across participants). After a 40-minute break, a BioSemi ActiveTwo system (BioSemi Inc., Amsterdam, The Netherlands) EEG cap and electrodes were fitted. Drug plasma levels have been found to reach maximum concentration after approximately 1.5-3h (Persiani, Rocchetti, Pacciarini, Holt, Toon, & Strolin-Benedetti, 1996; Agúndez, Garcia-Martin, Alonso-Navarro, & Jiménez-Jiménez, 2013). Approximately 1 hour and 20 minutes after drug intake, participants completed 6 minutes of sEBR recordings, followed by the backward masking task around 1.5h after drug intake (see below), during which EEG was recorded. After this task, the EEG setup was removed from the participant’s head. After a 30-minute lunch, participants proceeded to the attentional blink task, a simple reaction time task, and the probabilistic discrimination task (see below). At the end of the experiment, one final blood pressure/heart rate and VAS measure was undertaken. At the end of the final session, participants were asked to indicate in which session they believed they had received cabergoline.

### Physiological and subjective state measures

#### Heart rate and blood pressure

Physiological measurements were taken once during screening, and three times during both sessions; namely, on arrival, at around 1.5h after drug intake, and on completion of testing (± 3.5h after drug intake) (see Table 1). These measurements were obtained using an Omron® M3 comfort Sphygmomanometer.

#### Subjective self-report

A set of sixteen VAS measures were used (Bond & Lader, 1974), to assess the subjective state of the subject before medication intake, at around 1.5h after drug intake, and on completion of testing (± 3.5h after drug intake). Each scale consisted of a 100-mm horizontal line, anchored by contrasting states of mind (e.g., happy versus sad). Subjects were asked to regard each line as a continuum and to rate their feelings at the time by moving a vertical slider across each line. The scales could then be scored by measuring the length in millimeters from the positive end of each line to the subject’s marked location. These sixteen VAS measures were summarized as three categories: contentedness, calmness, and alertness (Bond & Lader, 1974).

#### Baseline dopamine proxies

Both sEBR and OSPAN are widely-used measures that have been related to baseline dopamine levels (Cools & D’Esposito, 2011; Jongkees & Colzato, 2016). Both measures have been used in combination with cabergoline in order to account for individual differences in baseline dopamine (Broadway et al., 2018; Cavanagh et al., 2014). Eye blink rate is defined as the number of spontaneous eye blinks per minute. The measure has high test-retest reliability (Kruis, Slagter, Bachhuber, Davidson, & Lutz, 2016) and is an often-used biomarker of baseline dopamine D2 receptor functioning (Jongkees & Colzato, 2016; Karson, 1983; Taylor et al., 1999, but see Sescousse et al., 2018). Subjects were asked to look at a central fixation cross on a computer screen in a relaxed state for 6 minutes while we measured eye activity from a set of vertical and horizontal electrodes, in order to detect eye blinks. This procedure was employed during all three lab visits.

Eye blinks were established in two ways. First, through a fully automatic procedure implemented in the python module MNE (*create_eog_epochs*; Gramfort et al., 2013). Second, eye-blinks were established through a custom semi-automatic procedure using EEGLAB for MATLAB (Delorme & Makeig, 2004). If mean sEBR per minute differed more than 3 blinks between both methods, the semi-automatic procedure was repeated, and an average was taken of both semi-automatic attempts as the final value. Prior to any repetition of the semi-automatic method, correlations between the automatic and semi-automatic method exceeded .95 for measurements during all three lab visits.

OSPAN is a working-memory task with high test-retest reliability (Unsworth et al., 2005), in which participants are instructed to remember letters, while solving simple arithmetic problems in between letter presentation (Unsworth et al., 2005). Sets of 3-7 letters were presented successively at fixation. The OSPAN score was calculated through partial credit scoring, so that each correctly recalled letter in the appropriate location was counted as correct, regardless of whether the entire sequence was recalled correctly or not. Scores could range from zero to 75. OSPAN was measured only during screening.

#### Reaction time

To assess the effects of cabergoline on alertness we administered a 40-trial simple reaction time (RT) task (Brown et al., 2016). In this task, participants had to respond as quickly as possible by pressing the spacebar whenever a white circle (subtending approximately 3.1° of visual angle) appeared at the center of the computer screen against a black background. Stimulus onset asynchrony was jittered between 500 and 1250 ms, with a mean of 1000 ms. This task lasted less than 2 minutes.

### Main experimental paradigms

All stimuli were presented on an ASUS VG236H 23-inch LCD screen (refresh rate = 100 Hz, resolution 1920×1080). Participants viewed the screen at a distance of 80 cm.

#### Backward masking

In the backward masking task, adapted from Van Opstal et al. (2014), participants had to indicate whether briefly presented masked digits (1, 4, 6, or 9) were smaller or larger than 5 and rate the confidence in their response (Figure 1). Each trial started with the presentation of a central fixation cross (30 point Courier New), which increased in size (106 point Courier New, 150 ms duration), cueing the impending target. The target stimulus (30 point Courier New) then appeared for 10 ms at one of two positions centered at the vertical midline (top or bottom, 2.29° from fixation). Both stimulus locations were equally probable. A mask followed the target (200 ms duration) at a variable stimulus onset asynchrony (SOA). Due to the employed refresh rate of 100Hz, the SOA could vary from 10 ms to 100 ms in 10 ms steps. By making the delay between cue and target dependent on SOA, the delay between cue and mask was held constant at 800 ms. The mask (30 point Courier New) was composed of two letters “E” and two letters “M”, tightly surrounding the target location without superimposing or touching it. All stimuli were black and presented on a white background, using the Psychophysics toolbox for MATLAB (Brainard, 1997). The central fixation cross was visible throughout the experiment.

**Fig. 1.**
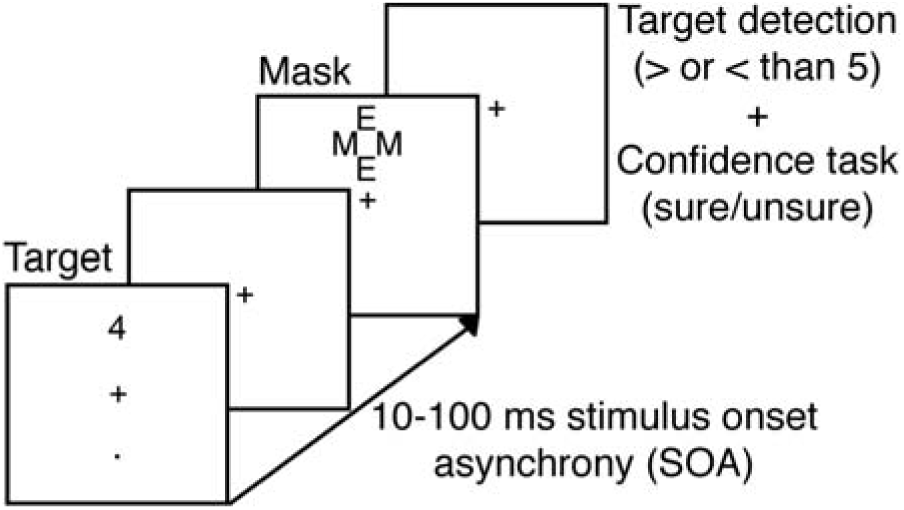
Experimental procedure for one trial of the backward masking task (adapted from Van Opstal et al., 2014)

Participants were instructed to indicate by button press whether the presented digit was smaller or larger than 5, while simultaneously indicating the confidence in their response (sure/unsure); resulting in four possible responses (<5 sure, <5 unsure, >5 unsure, >5 sure). In previous research it was found that the D2 agonist pergolide affected response confidence (Lou et al., 2011). Responses were given by means of a response box attached to the arm rests of the participant’s chair. Response buttons were counterbalanced across participants, who were instructed to guess one of two “unsure” buttons if they did not see the target.

If the participant’s reaction time exceeded 1 second, a message was presented indicating that their response was too slow for the duration of 1 second, urging a faster response. An individual threshold for awareness was established during the screening session (see above), by fitting a logistic model (threshold defined as SOA corresponding to 75% accuracy; mean threshold = 52.93 ms, min = 31 ms, max = 89 ms, sd = 14.38; Del Cul, Dehaene, & Leboyer, 2006; Van Opstal et al., 2014). This model was fitted on the basis of 176 trials during screening, where each of 11 SOA durations (from 0-100 ms) was presented 16 times.

Prior to the experiment, participants first completed a practice block (176 trials in screening, 88 trials during placebo and cabergoline sessions). In both drug sessions, participants completed 920 trials in total, split by seven possible SOAs between target and mask: 200 mask-only trials (0 ms SOA), 200 trials each for the main SOAs (10 ms / awareness threshold / 100 ms), 40 trials surrounding the threshold (threshold minus 10 ms and threshold plus 10 ms), and 40 trials with a 70 ms SOA. The trials in between individual thresholds and 100 ms were excluded from analysis (threshold + 10ms and 70 ms), because some participants arrived at an individual threshold at or above 70 ms. This meant that trials with a SOA at threshold + 10 ms and 70 ms would fall below or above participant’s individual threshold, depending on the participant. 80 trials per participant were discarded for this reason, leaving 840 trials.

#### Attentional blink

Participants also performed a standard AB task in which they had to identify two digits (T1 and T2) presented in a rapid stream of centrally presented distractors (letters and symbols; adapted from Slagter et al., 2012; Slagter et al., 2017; see Figure 2). T2 followed T1 either in the time window of the AB, after 200 ms (short-interval trial), or outside the time window of the AB, after 800 ms (long-interval trial). Each trial started with a central fixation cross (1500 ms), after which the stimulus stream began, consisting of 22 stimuli. Stimuli were presented on a black background (RGB 70, 70, 70) at the center of the screen (28 point Arial; 0.85° visual angle) for 50 ms, followed by a 50 ms blank. Digits were drawn randomly (without replacement) from the set 2–9. Distractors were randomly drawn (without replacement) from the following set of 30 letters and symbols: W, E, R, T, Y, U, P, A, D, F, G, H, J, K, L, Z, X, C, V, B, N, M, @, #, $, %, }, &, <, and =. Participants were asked to indicate sequentially the identity of the targets they saw, using the numpad on a standard keyboard. If they missed a target, they were instructed to guess. Stimulus presentation was performed using Presentation (Neurobehavioural Systems).

**Fig. 2.**
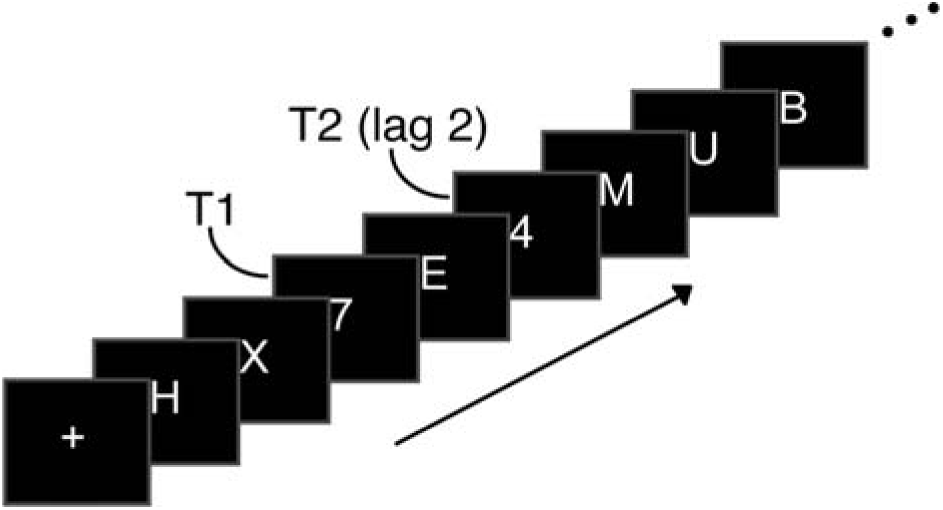
Experimental procedure for one trial of the attentional blink task (adapted from Slagter et al., 2017)

In both sessions, participants first completed a short practice block (20 trials), in which the first 8 trials moved at half speed. Next, participants moved on to the main experiment (222 trials), spread over 6 blocks consisting of 37 trials each.

#### Probabilistic discrimination

In the probabilistic discrimination task, adapted from Bauer et al. (2016, September), participants were presented continually with a central fixation cross (28 point Arial, RGB 0, 0, 0), on top of which an image (6.68° visual angle) of either a face or house was presented for 120 ms, against a grey background (RGB 128, 128, 128). Face stimuli were created on the basis of the Park Aging Mind Laboratory, University of Texas at Dallas (Minear & Park, 2004), while house stimuli were based on the Caltech University Computational Vision database (http://vision.caltech.edu/archive.html). On each trial, participants had to report the category of the image with by pressing “Q” or “P” on a standard keyboard. Responses were counterbalanced across participants, who were instructed to emphasize accuracy over and above speed. The maximum response interval was 1700 ms, after which the next stimulus was presented regardless of whether a response was given. The inter-trial interval was jittered and varied from 800 to 1200 ms.

The difficulty of stimulus discrimination was manipulated in terms of stimulus coherence (Figure 3). Stimuli from the above-mentioned databases were cropped to the outlines of faces and houses and a 2-dimensional spatial Fourier transform (on luminance values for x-/y-coordinates) was calculated. The amplitude (power-) spectra of all 82 face and house images were averaged and subsequently applied to all individual images, such that all images (faces and houses of all coherence levels) had an identical power spectrum. In other words, none of them differed in global contrast or luminance. The phase-spectra of each individual image (that therefore provided all pictorial information) was retained and was subsequently superimposed with various levels of (uniform) random noise for each image (Bauer et al., 2016, September). To account for bias in the circular phase-distribution of superimposed noise and signal phase-spectra, noise-spectra were sampled following previous suggestions (Dakin, Hess, Ledgeway, & Achtman, 2002).

**Fig. 3.**
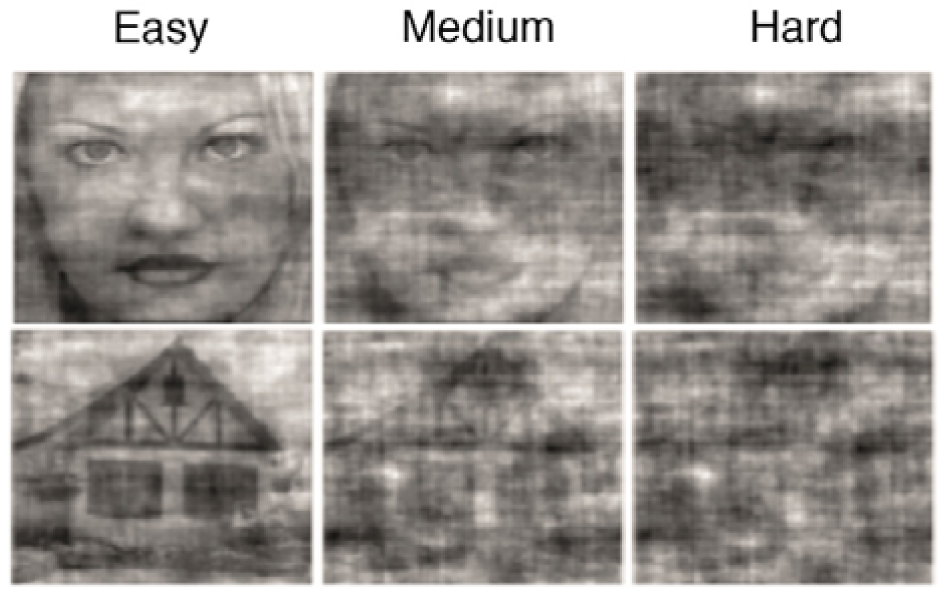
Probabilistic discrimination stimuli (adapted from Bauer et al., 2016, September)

Titration consisted of two subsequent procedures, in order to establish three difficulty levels for each individual participant. First, participants completed 300 trials, spread over 10 staircase blocks, in order to establish difficulty levels corresponding to an accuracy of 75% for each stimulus category separately. During this 3-up-1-down staircase procedure, participants received a green thumbs-up (RGB R [56 154 79], border RGB [17 79 22]) or red thumbs-down (RGB [83 2 5], border RGB [251 84 84]) at the center of the screen as feedback after each trial (500 ms duration).

Next, a total of 810 trials followed, across 27 blocks, in order to extrapolate the acquired difficulty level to three difficulty levels. For the second part of the titration procedure, participants received feedback in the break in between blocks; in order to counteract the development of a bias for one of two response categories. Difficulty levels were estimated using the method of constant stimuli (MOCS; Bauer et al., 2016, September). The psychometric functions obtained through this procedure were used to estimate difficulty (coherence) levels corresponding to 70, 82, and 95% accuracy.

In both the placebo and cabergoline session, the same three difficulty levels were employed that were acquired from the titration procedure during screening. Unbeknownst to participants, the prior probability of each category was manipulated in a block-wise manner (20/35/50/65/80%), spread over 25 blocks of 40 trials each, for a total of 1000 trials per session. As such, we manipulated perceptual information (difficulty) and stimulus prior probability independently of one another. Stimulus presentation for the titration procedure during the screening was performed using the Psychophysics toolbox for MATLAB (Brainard, 1997), and Presentation was used to present the task in both experimental sessions (Neurobehavioural Systems).

### Behavioral analyses

#### Physiological and subjective state measures

In order to test whether physiology and subjective state changed over the course of the experiment, and whether these measures were influenced by cabergoline, we conducted a repeated-measures analysis-of-variance (RM ANOVA) for heart rate, diastolic and systolic blood pressure, and each VAS category separately; across all three time-points and both sessions. We conducted a paired-samples t-test between sessions for our simple RT task, as an additional measure to assess alertness. In order to test whether cabergoline exerted influence on sEBR, we conducted a paired-samples t-test between sEBR under placebo versus cabergoline. Kendall’s Tau correlation was employed to establish the relationship between sEBR sessions, as well as the relationship between sEBR and OSPAN, as this coefficient is more robust in the case of small samples and tied ranks (Bonett & Wright, 2000). Correlations were Bonferroni corrected for multiple comparisons.

#### Backward masking

The dependent measures in the backward masking task were accuracy (0/1) and confidence (unsure/sure). For both of these measures, we computed a 2 (Drug; placebo/cabergoline) x 5 (SOA; mask-only/10 ms/threshold – 10 ms/threshold/100 ms) RM ANOVA. Furthermore, we performed an additional analysis including screening sEBR and OSPAN as covariates in both of these analyses, as we predicted cabergoline effects may depend on individual baseline dopamine levels, based on previous reports (Broadway et al., 2018; Cavanagh et al., 2014; Cools & D’Esposito, 2011; Jongkees & Colzato, 2016). We repeated these analyses for the two-alternative forced choice version of the signal detection theory parameters d’ (Green & Swets, 1966) and meta-d’ (Maniscalco & Lau, 2012), for the three primary SOAs (10 ms, threshold, and 100 ms).

One participant confused the confidence response buttons in one session. We reversed these confidence scores manually. One participant experienced side effects only near the end of the last session. This participant thus completed the backward masking task twice without knowledge about drug condition. In order to maximize statistical power, this participant was included in all analyses concerning the backward masking task (including EEG), but not in the analyses of other tasks. It did not matter for our results whether this participant was included in the backward masking task analyses or not.

#### Attentional Blink

The dependent measures for the AB task were T1 accuracy and T2 | T1 accuracy. In other words, T2 accuracy was based only on those trials where T1 was correctly reported. For each of these measures, we computed a 2 (Drug; placebo/cabergoline) x 2 (Lag; 2/8) RM ANOVA. For this task as well, we computed additional analyses in order to include sEBR and OSPAN as covariates. Finally, we computed AB size, in order to investigate the relationship between AB size and our baseline dopamine measures as some (Colzato, Slagter, Spapé, & Hommel, 2008) but not other studies (Slagter et al., 2012) have found.

#### Probabilistic discrimination

In the case of the probabilistic discrimination task, our dependent measure of interest was accuracy. As such, we computed a 2 (Drug; placebo/cabergoline) x 3 (Difficulty; easy/medium/hard) x 5 (Probability; .2/.35/.5/.65/.8) RM ANOVA. For this analysis as well, we computed additional models including screening sEBR and OSPAN as covariates.

Our titration procedure was not successful for all participants. As a result, a number of participants ended up with only two difficulty levels for one out of two stimuli. For this reason, we repeated the above analysis for both the group with all difficulty levels (N = 16), and participants who had either two or three difficulty levels as a result of the titration procedure (N = 24). In this latter analysis, we excluded the medium difficulty trials. One participant was excluded from both analyses, because this participant ended up with only one difficulty level for face stimuli.

All covariates in the above-mentioned RM ANOVAs were centered (van Breukelen & van Dijk, 2007). For all repeated-measures ANOVA analyses, whenever Mauchly’s test suggested a violation of sphericity, we report Geenhouse–Geisser corrected *P*-values, but uncorrected degrees of freedom. In order to test for order effects, we repeated each of the RM ANOVAs for our behavioral paradigms including a between-subject factor indicating whether a participant received either placebo or cabergoline in the first session.

#### Bayesian statistics

In order to evaluate evidence in favor of our (null) hypotheses, we conducted Bayesian statistics. For each reported frequentist test, we report the Bayes factor corresponding to the inclusion of a factor or interaction within the model in question (shortened to BF_incl_), compared to equivalent models stripped of the effect. For example, BF_incl_ = 10 indicates that a model including the factor in question is ten times more likely given the data compared to a model without the variable. Conversely, BF_incl_ = .1 indicates that a model without said effect is ten times more likely given the data. All Bayesian statistics were conducted using JASP (2019, version 0.10.0).

All data visualization was performed with the help of raincloud plots (Allen, Poggiali, Whitaker, Marshall, & Kievit, 2019), which include the mean, individual data points, as well as the overall distribution of the measure in question.

### EEG

#### Recording and preprocessing

EEG data, digitized at 512 Hz, were continuously recorded in both the placebo and cabergoline session during 6 minutes of sEBR and the backward masking task, using an ActiveTwo system (BioSemi, Amsterdam, the Netherlands), from 64 scalp electrodes placed according to the 10/20 system, four electro-oculographic electrodes placed above and below, and to the side of the eyes, and two external electrodes attached to each earlobe. EEG data were offline referenced to the average activity recorded at the earlobes, and high-pass ‘firws’ filtered (default settings) at 0.05 Hz using a Kaiser window, following previous suggestions (Widmann, Schröger, & Maess, 2015). The continuous data were subsequently epoched from −1.5 to 1.5 s around stimulus presentation and baseline corrected to the average activity between −200 ms and 0 ms pre-stimulus. Epochs containing EMG artifacts or eye blinks surrounding stimulus presentation were rejected based on visual inspection. Extremely noisy or broken channels were interpolated. Remaining eye blink artifacts were removed by decomposing the EEG data into independent sources of brain activity using an Independent Component Analysis, and removing eye blink components from the data for each subject individually. Epochs were low-pass filtered at 30 Hz for visualization purposes only. Preprocessing was done using the Fieldtrip toolbox (Oostenveld, Fries, Maris, & Schoffelen, 2011) for Matlab (The MathWorks, Inc. Natick, MA, USA) using custom-written Matlab scripts.

#### Analyses

To determine the effect of our manipulations on ERP markers of information-processing, we examined the effects of SOA and Drug on the amplitude of the visual-evoked P1 and N1 components, the N2, as well as of the later P3b (Del Cul, Baillet, & Dehaene, 2007). In line with Del Cul et al (2007), we epoched the ERP data to the onset of the mask. Next, we subtracted the data from the mask-only SOA condition from all other SOA conditions. Finally, we shifted ERP onset back to target onset, in order to compute target-locked ERPs. Visual inspection of the grand- and condition-average ERPs showed that the P1 and N1 components peaked over lateral occipitoparietal scalp sites (PO7, PO3, O1, PO4, PO8, O2), the N2 over centroparietal scalp regions (C1, Cz, C2, CP1, CPz, CP2), and the P3b over central parietal scalp sites (P1, Pz, P2, PO3, POz, PO4). These scalp sites were used to determine the peak amplitude and latency of these components for each condition of interest. Specifically, the largest positive voltage value between 75–150 ms post-target, and the largest voltage negativity within 150–225 ms were selected to determine the amplitude and latency of the P1 and N1 peaks, respectively, for each subject separately. In the case of the P2 and N2, these intervals were 175-250 ms, and 250-375 ms, respectively. For the P3, an interval of 300-450 ms was used. All average amplitude values 15 ms around the peak sample, as well as individual latencies were entered into separate RM ANOVAs with two within-subject factors: SOA (10 ms, threshold-10 ms, threshold, 100 ms), and Drug (placebo/cabergoline). Because the mask-only condition is used to acquire ERP data for the remaining four SOA conditions (see above), the SOA factor contains four instead of five levels for ERP analyses.

## Results

### Physiological and subjective state measures

First, we aimed to establish whether cabergoline exerted physiological effects (Figure 4). Heart rate decreased over time in both placebo and cabergoline conditions (F_(2, 48)_ = 11.8, p < .001, η^2^ = .06, BF_incl_ > 100), and was overall higher in the cabergoline condition (F_(1, 24)_ = 9, p = .006, η^2^ = .03, BF_incl_ = 57.3), but there was no interaction between Time and Drug (F_(2, 48)_ < 1, p = .63, BF_incl_ = .14; see Figure 4a). In the case of diastolic blood pressure as well, we also observed a decrease over time (F_(2, 48)_ = 21.7, p < .001, η^2^ = .14, BF_incl_ > 100), but in this case an overall lower measurement in the cabergoline condition (F_(1, 24)_ = 9.9, p = .004, η^2^ = .03, BF_incl_ = 12.4), and again no interaction between Time and Drug (F_(2, 48)_ < 1, p = .42, BF_incl_ = .2). With regards to systolic blood pressure, we also observed a decrease in blood pressure over time (F_(2, 48)_ = 36.7, p < .001, η^2^ = .22, BF_incl_ > 100) and an overall lower measurement in the cabergoline condition (F_(1, 24)_ = 9, p = .006, η^2^ = .03, BF_incl_ = 31.2), but here a significant interaction between Time and Drug was found (F_(2, 48)_ = 7.2, p = .002, BF_incl_ = 5.8; see Figure 4b). Thus, all three physiological measures decreased over time. While heart rate was higher, and diastolic blood pressure lower across the cabergoline session, cabergoline can only be said to have decreased systolic blood pressure, as indicated by the interaction between Time and Drug.

**Fig. 4.**
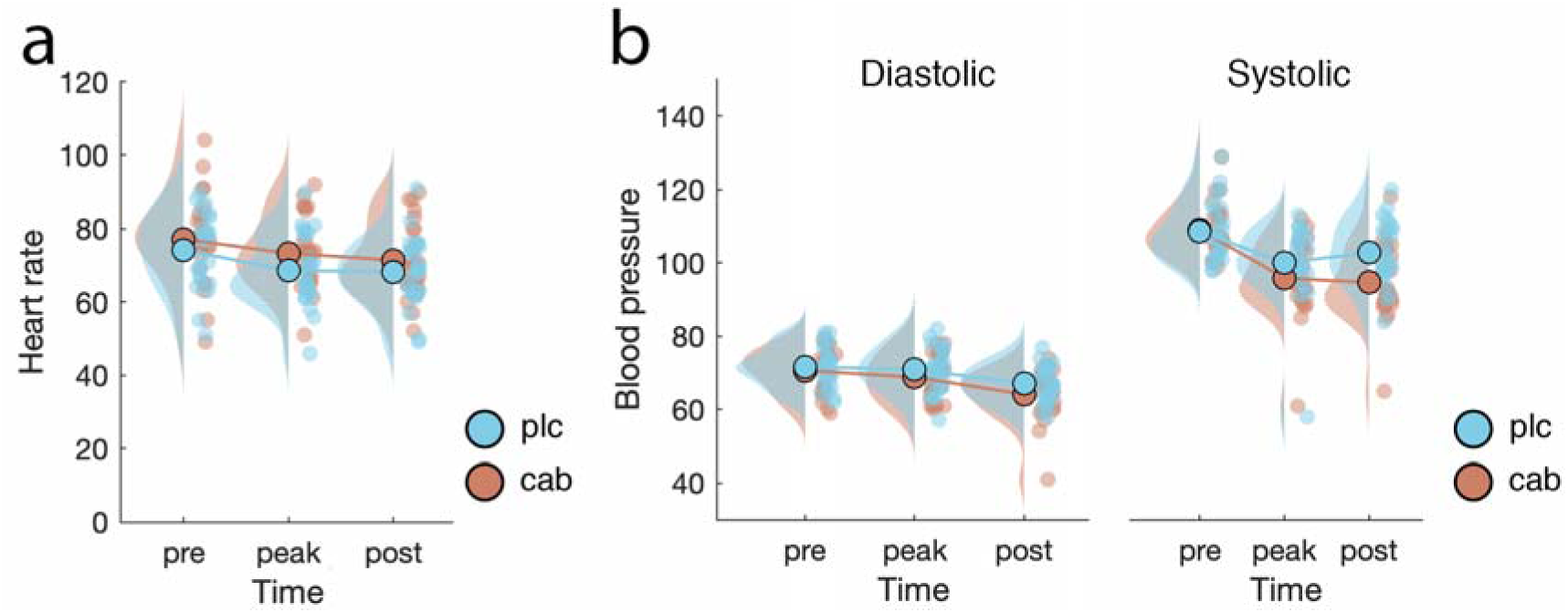
Time course and difference between drug conditions in heart rate and blood pressure. All three physiological measures decreased over time. Cabergoline only affected systolic blood pressure, while heart rate was higher, and diastolic pressure was lower in the cabergoline versus placebo session

Cabergoline also affected the subjective state of participants over the course of the experiment, as indicated by the VAS (visual analog scale, see Figure 5). On the calmness scale, there was no main effect of Drug (F_(1, 24)_ < 1, p = .63, BF_incl_ = .23), but calmness increased over the course of the experiment (F_(2, 48)_ = 14.5, p < .001, η^2^ = .04, BF_incl_ = 31). In addition, we found an interaction between Drug and Time (F_(2, 48)_ = 5.8, p = .005, η^2^ = .02, BF_incl_ = 2), reflecting the fact that calmness ratings increased less in the cabergoline condition (Figure 5). In the case of contentedness, there was no main effect of Drug (F_(1, 24)_ < 1, p = .46, BF_incl_ = .3), Time (F_(2, 48)_ = 2.7, p = .08, BF_incl_ = .33), or an interaction between Drug and Time (F_(2, 48)_ < 1, p = .53, BF_incl_ = .16; see Figure 5). Finally, alertness decreased over the course of the experiment (F_(2, 48)_ = 10.3, p = .001, η^2^ = .12, BF_incl_ > 100) and was generally lower in the cabergoline session compared to placebo session (F_(1, 24)_ = 6.3, p = .02, η^2^ = .01, BF_incl_ = 1.1), but there was no interaction (F_(2, 48)_ = 1.8, p = .18, BF_incl_ = .21; see Figure 5). Thus, cabergoline decreased self-reported ratings of calmness and alertness as the experiment progressed.

85% of participants successfully guessed when they received cabergoline at the end of the experiment. However, despite these effects, we found no significant difference in sEBR (spontaneous eye blink rate) between placebo (M = 14 blinks per minute, SD = 8.9) and cabergoline (M = 15.3, SD = 10.6) (t_(24)_ = −1.2, p = .23, BF_01_ = .41). In addition, we were unable to replicate the finding by Cavanagh and colleagues (2014) that baseline sEBR conditioned a cabergoline-induced shift in line with an inverted-U-shape pattern, to the point where high baseline sEBR (as indicative of high tonic striatal dopamine) is associated with a reduction in blink rate, whereas low baseline sEBR is linked to an increase in blink rate: we found that the difference in sEBR between drug conditions was unrelated to screening sEBR (M = 13.2, SD = 7.8) (*τ*_(25)_ = .23, p = .11).

**Fig. 5.**
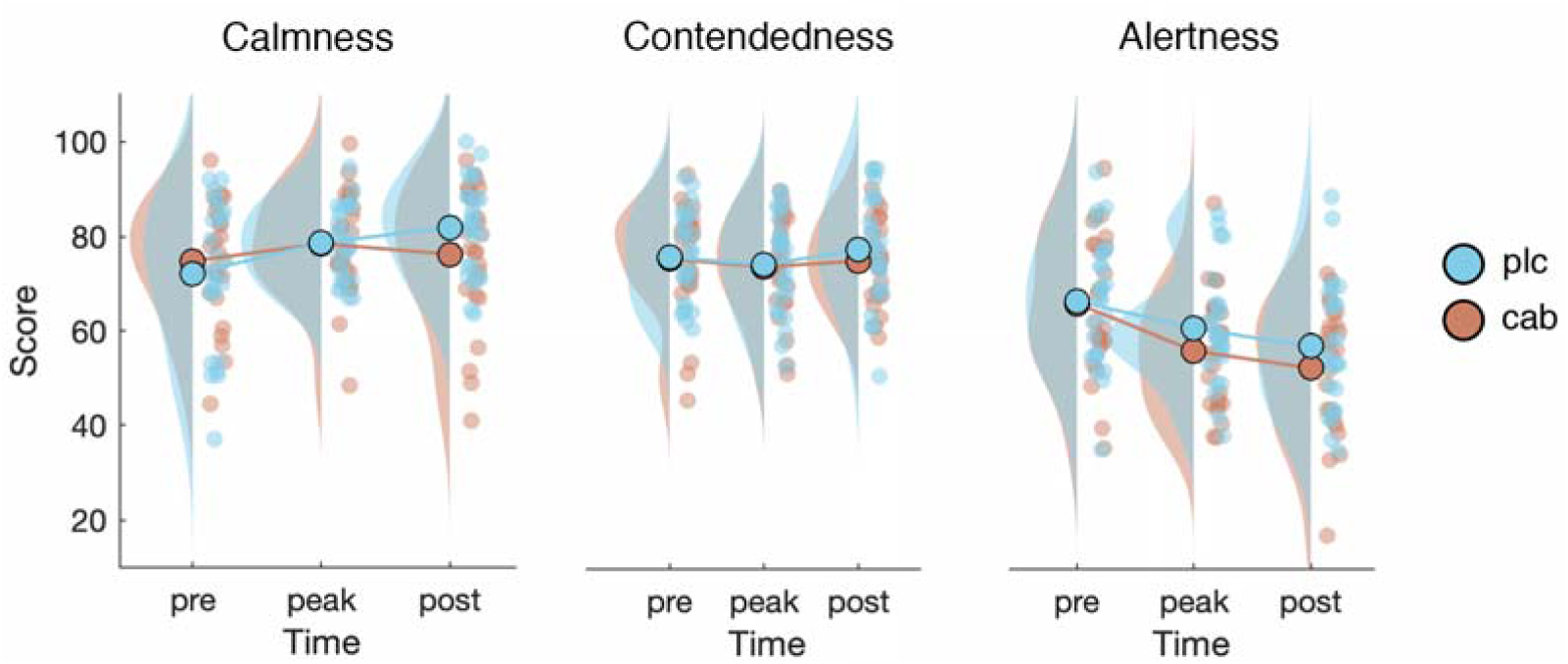
Time course and difference between drug conditions in VAS scores for calmness, contentedness, and alertness. Participants become calmer and less alert over the course of the experiment. Cabergoline stifled this increase in calmness

Replicating previous findings (Jongkees & Colzato, 2016; Kruis et al., 2016), sEBR was correlated across sessions: between the placebo and cabergoline session (*τ*_(25)_ = .63, p < .001), between the screening and cabergoline session (*τ*_(25)_ = .37, p = .01), as well as between the screening and placebo session (*τ*_(25)_ = .32, p = .03; see Figure 6a). The latter two correlations did not survive a Bonferroni correction for multiple comparisons using a corrected alpha level of .05 / 6 = .0083, based on all correlations computed in this section. Together these results lend support to the robustness of the measure when measured at the same time of day (Barbato et al., 2000; Jongkees & Colzato, 2016). However, despite the proposed relation between sEBR and OSPAN as a biomarker of striatal dopamine, we found no relationship between these measures in either the cabergoline (*τ*_(25)_ = .06, p = .67) or placebo session (*τ*_(25)_ = .06, p = .67). This relationship was absent even when these measures were collected in the same session; namely, during screening (*τ*_(25)_ = -.11, p = .47; see Figure 6b). Finally, alertness, as indicated by our simple RT task also did not differ between the placebo (M = 224 ms, SD = 15.8) and cabergoline condition (M = 225 ms, SD = 15.1) (t_(25)_ = -.54, p = .59).

**Fig. 6.**
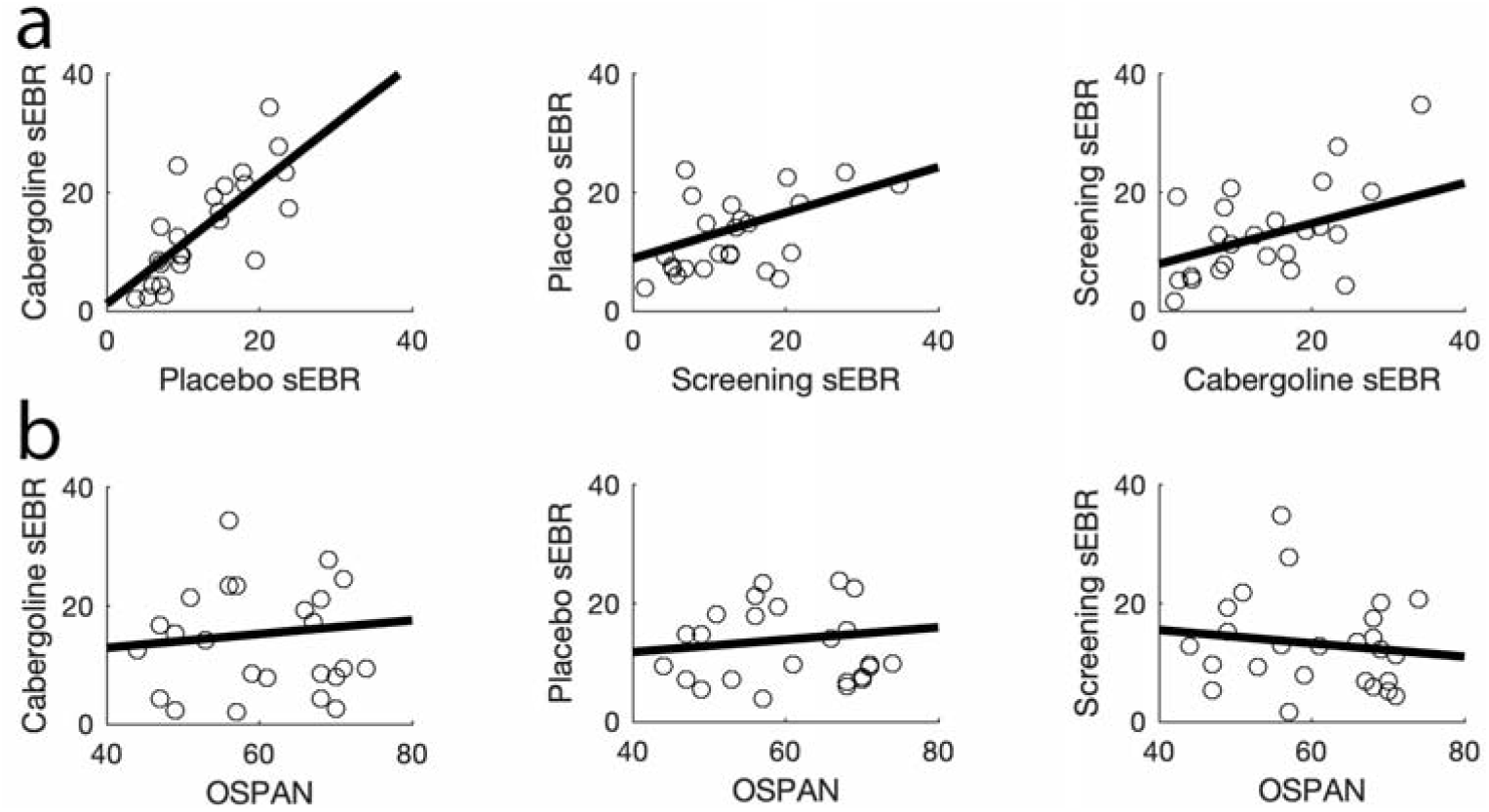
The relationship between sEBR measures among themselves and in relation to OSPAN. All sEBR measures were found to be positively correlated (**a**), but we found no relationship between sEBR and OSPAN (**b**)

Thus, cabergoline did not affect sEBR or our objective measure of alertness.

### Main experimental paradigms

We next examined potential effects of cabergoline on our main experimental measures of interest: target identification accuracy and processing in the backward masking task, attentional blink size in the attentional blink task, and discrimination accuracy under varying conditions of difficulty and probability in the probabilistic discrimination task. To foreshadow our results, cabergoline did not affect any of the key behavioral findings associated with these tasks, whether sEBR or OSPAN were included as covariates in the analyses or not. Neither did it affect neural processing of the target in the backward masking task, as shown by ERP analyses. Yet, importantly, we did replicate all standard findings typically obtained with these tasks (e.g., effects of masking on target-evoked ERPs, the attentional blink).

### Backward masking

#### Behavior

As is typically observed (e.g., Breitmeyer, 2007) and shown in Figure 7, targets were more often identified correctly in the backward masking task as the delay between target and mask (SOA) increased (F_(4, 100)_ = 187.4, p < .001, η^2^ = .79, BF_incl_ > 100). Yet, in contrast to our main prediction that cabergoline would affect participant’s ability to detect targets, there was no interaction between Drug and SOA (F_(4, 100)_ < 1, p = .56; BF_incl_ = .03; see Figure 7). We also did not find an overall difference between placebo and cabergoline on target identification accuracy (F_(1, 25)_ < 1, p = .85, BF_incl_ = .13). Controlling for screening sEBR, OSPAN, or the combination of both did not change the results. Neither did the inclusion of a between-subject factor for drug order. We repeated these analyses with d’ as the dependent variable, but results were equivalent.

**Fig. 7.**
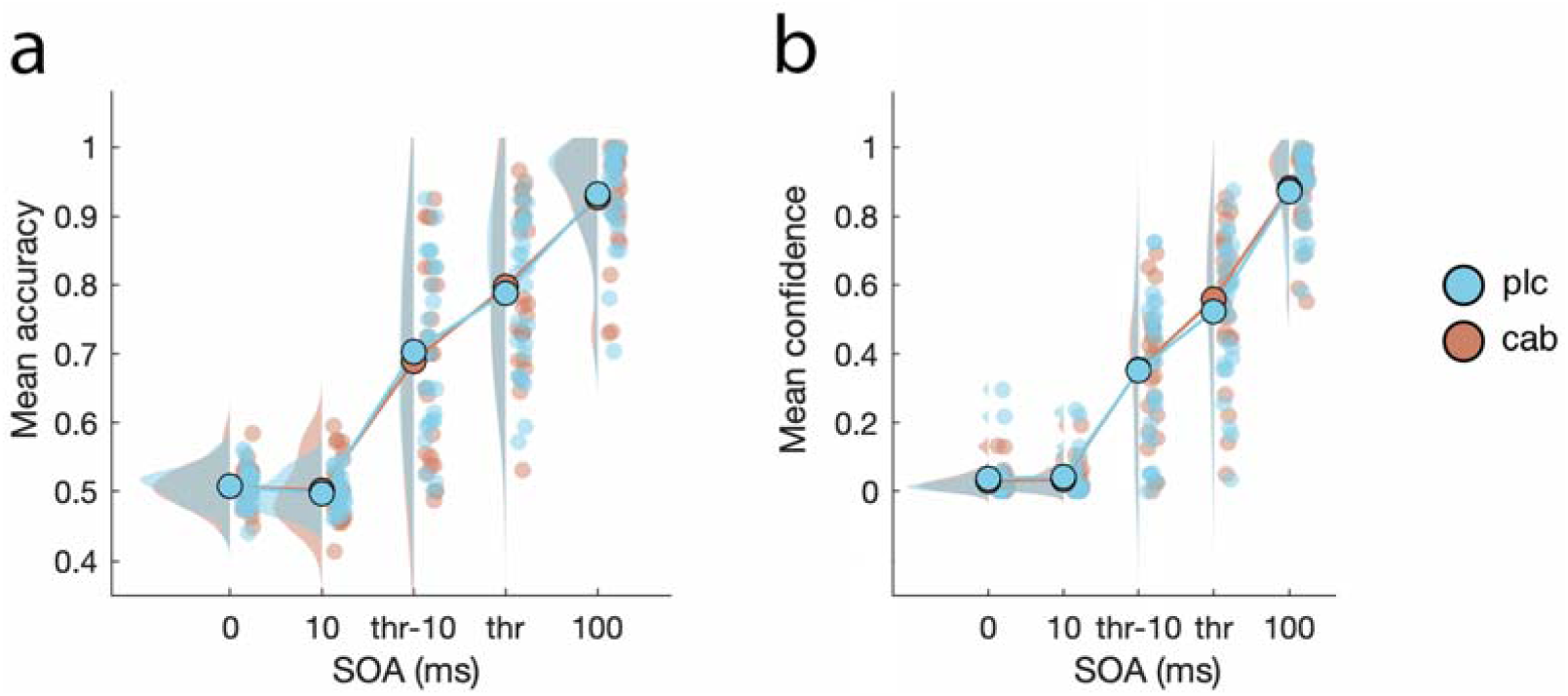
Mean accuracy and confidence scores in the backward masking task across five possible SOAs and both drug conditions. While both accuracy (**a**) and confidence (**b**) scaled with the duration of SOA, cabergoline had no effect on either of these measures

Similarly, in the case of confidence scores, cabergoline did not affect reported confidence ratings, as indicated by the lack of an interaction between SOA and Drug (F_(4, 100)_ = 1.6, p = .21; BF_incl_ = .06; see Figure 7). We also found no overall difference between drug conditions (F_(1, 25)_ < 1, p = .63; BF_incl_ = .14), but participants did report an improvement in response confidence as SOA increased (F_(4, 100)_ = 204.8, p < .001, η^2^ = .8; BF_incl_ > 100). Again, correcting for baseline dopamine measures or drug order had no impact on the results. As with d’, repeating these analyses with meta-d’ as the dependent variable made no difference. Thus, we replicated a similar pattern of results with regards to both objective and subjective aspects of the participant’s response during backward masking (Del Cul et al., 2007), but these patterns were not affected by cabergoline.

#### EEG

In addition to these behavioral masking effects, we replicated previous reports that target-evoked ERP components scale with the duration of target-mask SOA (Del Cul et al., 2007). We found a strong effect of SOA on the amplitude and peak-latency of the target-evoked P1, N1, N2, and P3b components. However, in the case of our ERP results as well, each of these measures was unaffected by cabergoline (see Table 2 and Figure 8). In order to remain close to previous studies (Del Cul et al., 2007), we repeated these analyses with an average reference instead of an earlobe reference, but the conclusions belonging to these results remained the same. Thus, cabergoline did not affect the threshold for conscious perception by modulating neural target processing.

**Table 2.** Summary of the statistical analyses performed on ERP data.

| Component | Measure | SOA ($F_{(3, 75)} =$) | Drug ($F_{(1, 25)} =$) | SOA*Drug ($F_{(3, 75)} =$) |
| --- | --- | --- | --- | --- |
| P1 | Amplitude | 51.7, $p < .001^*$ | .01, $p = .91$, $BF_{incl} = .16$ | .62, $p = .60$, $BF_{incl} = .08$ |
| | Latency | 8.2, $p < .001^*$ | .17, $p = .68$, $BF_{incl} = .16$ | .48, $p = .7$, $BF_{incl} = .08$ |
| N1 | Amplitude | 12.4, $p < .001^*$ | .04, $p = .85$, $BF_{incl} = .15$ | .44, $p = .73$, $BF_{incl} = .06$ |
| | Latency | 66.8, $p < .001^*$ | .33, $p = .57$, $BF_{incl} = .19$ | .07, $p = .98$, $BF_{incl} = .06$ |
| N2 | Amplitude | 4.7, $p = .005^*$ | .04, $p = .83$, $BF_{incl} = .15$ | .21, $p = .89$, $BF_{incl} = .06$ |
| | Latency | 99.3, $p < .001^*$ | 2.9, $p = .1$, $BF_{incl} = .38$ | .09, $p = .97$, $BF_{incl} = .06$ |
| P3b | Amplitude | 57.2, $p < .001^*$ | .02, $p = .9$, $BF_{incl} = .15$ | .9, $p = .44$, $BF_{incl} = .07$ |
| | Latency | 10, $p < .001^*$ | .1, $p = .74$, $BF_{incl} = .15$ | 1.7, $p = .18$, $BF_{incl} = .09$ |

\* = $BF_{incl} > 100$

**Fig. 8.**
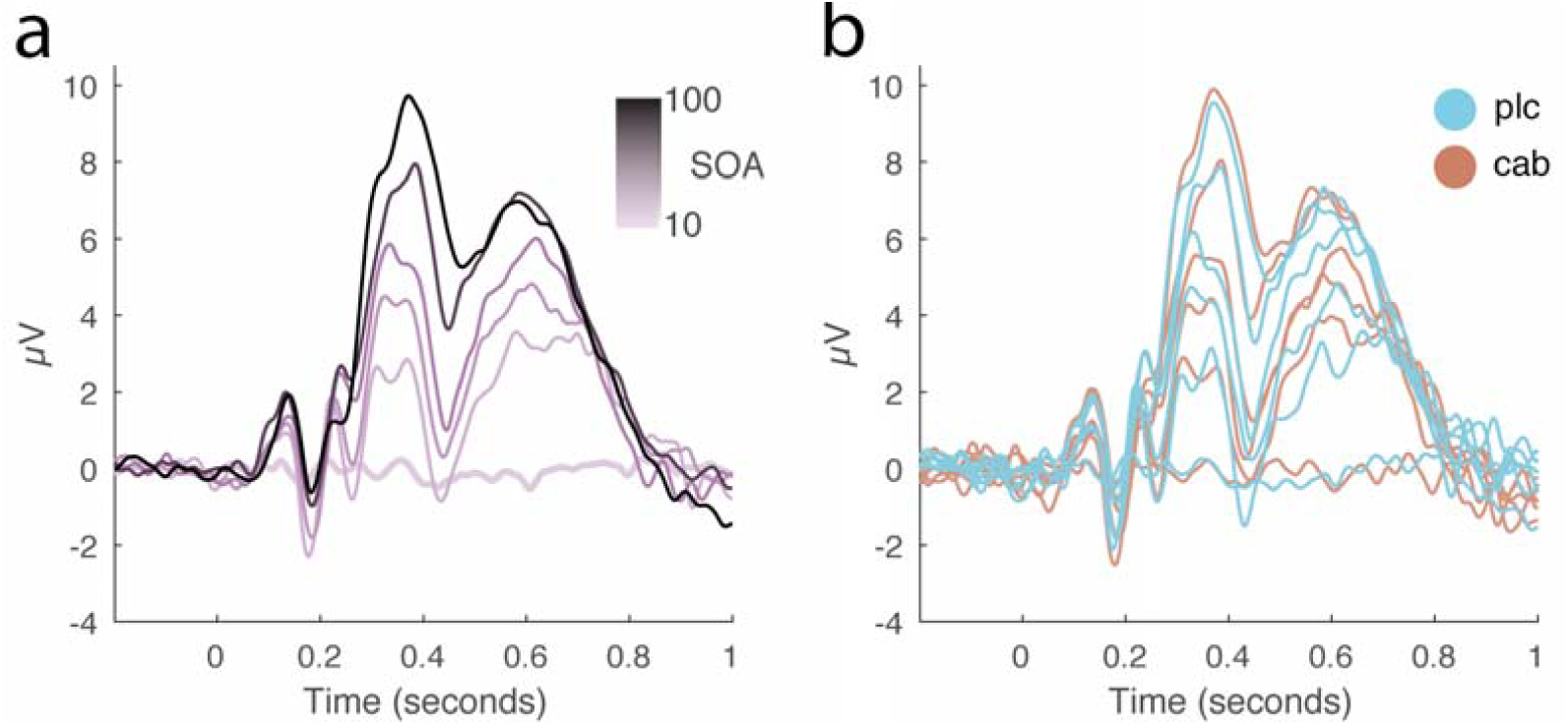
Effects of cabergoline on target-evoked ERPs in the backward masking task. This figure displays the grand-average target-evoked ERPs for P3b electrodes (P1, Pz, P2, PO3, POz, PO4) for both drug conditions combined (**a**) and separately (**b**), per target-mask SOA condition (10, 40, 50, 60, 70, and 100 ms). This figure shows that while ERP amplitudes and latencies generally increased as a function of target-mask SOA, these measures were unaffected by cabergoline.

### Attentional blink

As expected and shown in Figure 9, we found a robust attentional blink: T2|T1 accuracy was significantly worse when T2 was presented after one distractor (lag 2) compared to seven distractors (lag 8; F_(1, 24)_ = 43.9, p < .001, η^2^ = .39; BF_incl_ > 100). T2|T1 accuracy was marginally better in the cabergoline condition (81.9±11%) compared to placebo (79.7±14%; F_(1, 24)_ = 4.2, p = .052; BF_incl_ = .34), but we found no interaction between Lag and Drug (F_(1,_ 24) = 1.42, p = .25; BF_incl_ = .3).

**Fig. 9.**
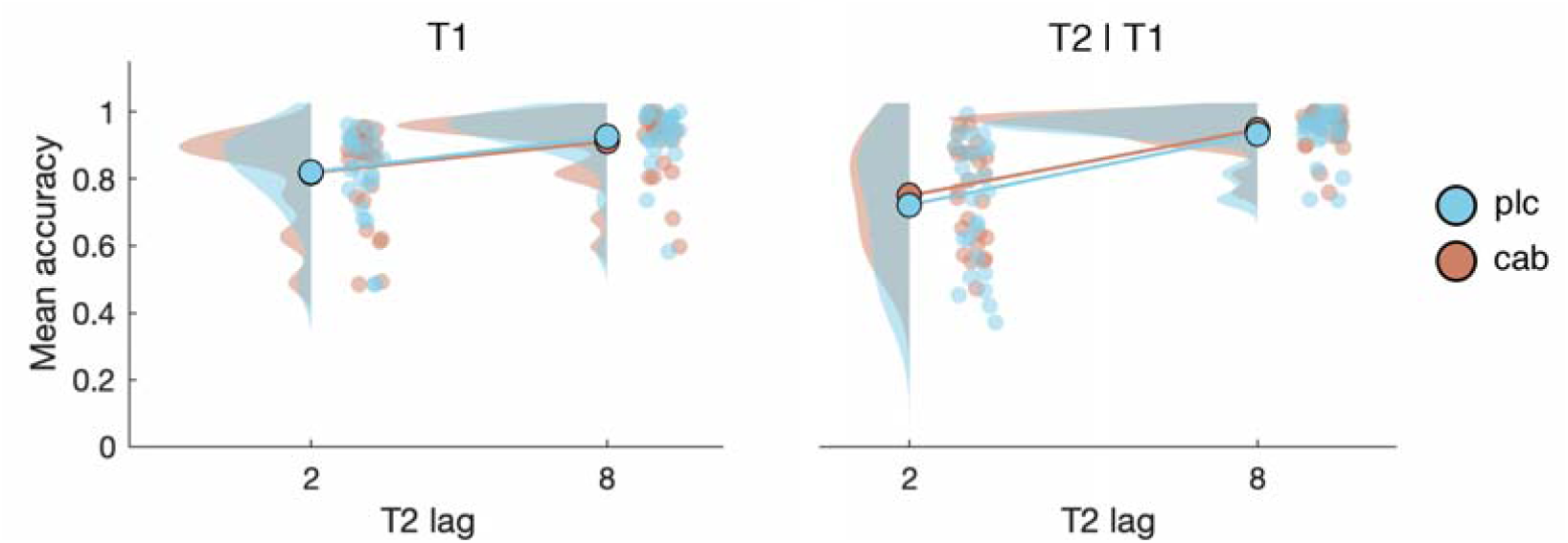
Mean accuracy for T1 and T2|T1 at lag 2 and lag 8. While we replicated behavioral findings common to the attentional blink paradigm, cabergoline did not affect the attentional blink

When we controlled for baseline dopamine measures (i.e., baseline OSPAN and sEBR), the inclusion of OSPAN did not affect the results, and controlling for sEBR also did not reveal a critical interaction between Lag and Drug (F_(1, 22)_ = 1.3, p = .26; BF_incl_ = .3). We did find a between-subject effect of screening sEBR (F_(1, 22)_ = 5.2, p = .03; BF_incl_ = 2.5), which exacerbated the main effect of Drug (F_(1, 22)_ = 5.4, p = .029; η^2^ = .004; BF_incl_ = .36). In addition, we found an interaction between sEBR and Drug (F_(1, 22)_ = 5.8, p = .025; η^2^ = .005): sEBR correlated with overall T2|T1 accuracy in the placebo (*τ*_(25)_ = -.31, p = .03), but slightly less so in the cabergoline (*τ*_(25)_ = -.26, p = .07) condition.

Although the critical three-way interaction between sEBR, Lag and Drug was not significant, we also found an interaction between screening sEBR and Lag (F_(1, 22)_ = 5, p = .036; η^2^ = .004). This interaction is best understood in terms of the relationship between sEBR and AB size (Colzato et al., 2008; Slagter et al., 2012); namely, the difference in T2|T1 accuracy between lag 2 and 8. AB size ranged from −2.9% to 59.7% in the placebo condition, and from 0% to 47.5% in the cabergoline condition. The interaction between sEBR and Lag stems from a small correlation between screening sEBR and AB size in the cabergoline (*τ*_(25)_ = .31, p = .03) and placebo condition (*τ*_(25)_ = .28, p = .047). We found no other relationships between dopamine baseline measures and the AB; the difference in AB size between the cabergoline and placebo session was unrelated to screening sEBR (*τ*_(25)_ = .05, p = .71). For the cabergoline session, there was no relation between AB size and sEBR (*τ*_(25)_ = .13, p = .35), nor between AB size and OSPAN (*τ*_(25)_ = .08, p = .57; see Figure 10a). Similarly, for the placebo session we did not find a relationship either between AB size and sEBR (*τ*_(25)_ = .15, p = .3), nor OSPAN (*τ*_(25)_ = .02, p = .89, see Figure 10b). However, none of the correlations reported in this section survived a Bonferroni correction for multiple comparisons using a corrected alpha level of .05 / 9 = .0056, based on all correlations computed in this section. Thus, we found no strong support for a relationship between sEBR and the AB, or OSPAN and the AB, nor did controlling for these factors reveal drug-related effects on the AB.

**Fig. 10.**
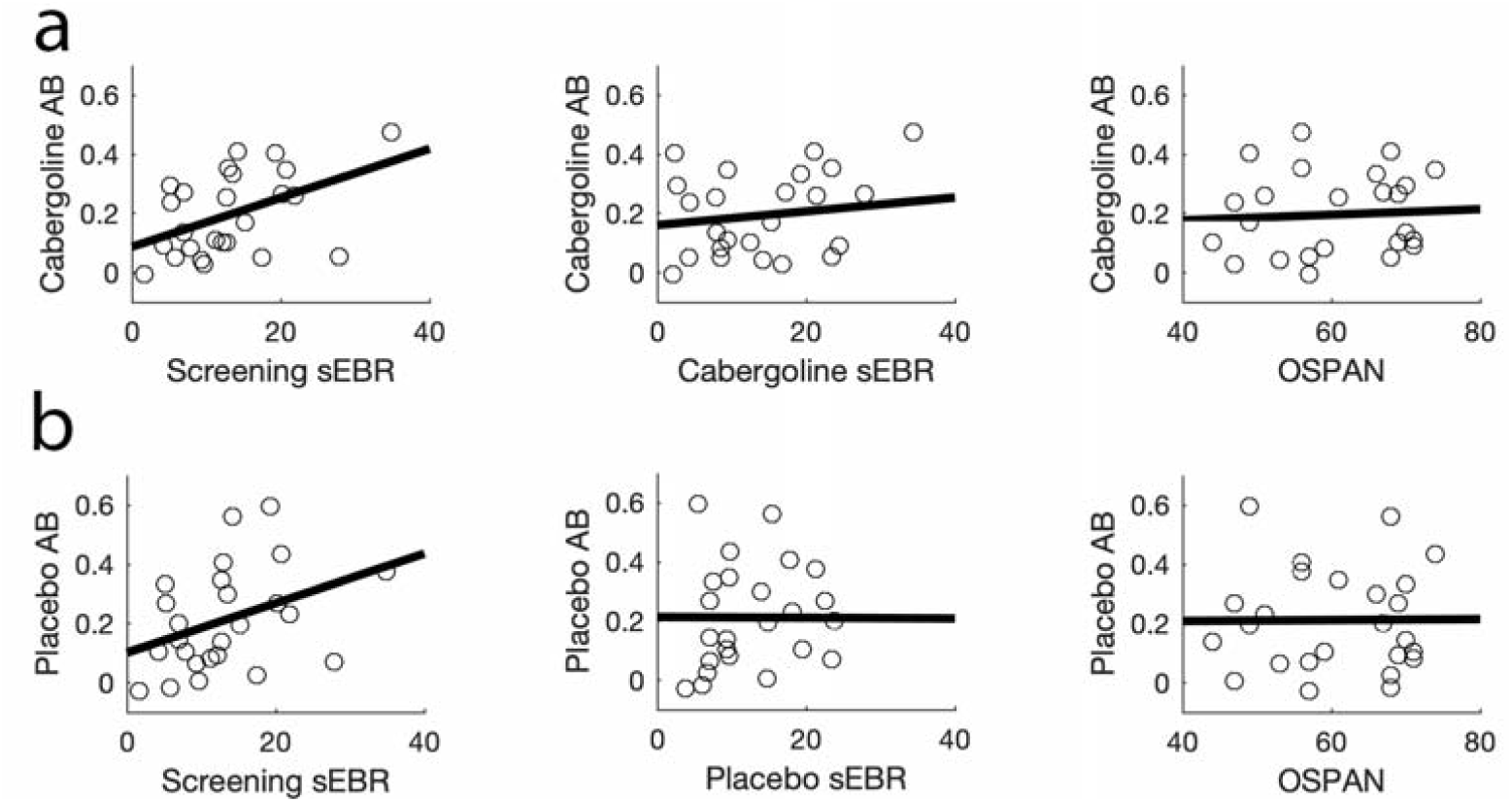
The relationship between AB size and striatal-dopamine proxy measures. Both for the cabergoline and placebo condition, we found a positive relation between AB size and screening sEBR, while such a relationship was not present neither for sEBR measured within the same session, nor for OSPAN

When controlling for drug order, we found an interaction between Drug and drug Order (F_(1, 21)_ = 5.7, p = .027; η^2^ = .004; BF_incl_ = .43). When participants received placebo first, they were less accurate overall in identifying T2|T1 in the placebo condition (77.5±15%), compared to the cabergoline condition (81.6±11%). For participants who received cabergoline first, this difference was not present (placebo: 82.1±12%, cabergoline: 82.3±10%). No other order effects were present. Together these results indicate that cabergoline did not affect the attentional blink.

We found no difference in T1 accuracy between the placebo and cabergoline condition (F_(1, 24)_ < 1, p = .49; BF_incl_ = .27), or an interaction between Drug and Lag (F_(1, 24)_ < 1, p = .5; BF_incl_ = .32), but T1 accuracy was worse when T2 was presented after only one distractor (lag 2; F_(1, 24)_ = 46.6, p < .001, η^2^ = .16; BF_incl_ > 100). Controlling for sEBR, OSPAN, or drug order did not affect the results for T1 accuracy. Thus, cabergoline also did not affect T1 identification.

### Probabilistic discrimination

In our final experimental paradigm, participants were tasked with discriminating between face and house stimuli that varied in difficulty and probability of occurrence (see Methods). Cabergoline did not affect discrimination accuracy (specifically: hit-rate; F_(1, 23)_ = 2.4, p = .13, BF_incl_ = .35). Neither did we find an interaction between Drug and Difficulty (F_(1, 23)_ < 1, p = .91, BF_incl_ = .14), Drug and Probability (F_(4, 92)_ < 1, p = .78, BF_incl_ = .02), or a three-way interaction (F_(4, 92)_ < 1, p = .48, BF_incl_ = .05; see Figure 11a). Thus, cabergoline did not significantly affect perception on this task either. Standard effects observed with this task in non-drug studies were replicated (Bauer et al., 2016, September): accuracy was lower on difficult (F_(1, 23)_ = 208.3, p < .001, η^2^ = .60, BF_incl_ > 100), and on low-probability trials (F_(4,_ 92) = 22.49, p < .001, η^2^ = .04, BF_incl_ > 100). When we controlled for screening sEBR and OSPAN, this did not change the results. We did find an interaction between drug Order and Drug (F_(4, 88)_ = 5.2, p = .03, BF_incl_ = 1.3). Opposite to what we found for the attentional blink task, when participants received placebo first, they were more accurate overall in the placebo (84±7%) compared to the cabergoline (82.4±7%) condition. When cabergoline was administered first, this difference was absent (placebo: 83.5±5%, cabergoline: 83.1±4%). We found no other order effects.

**Fig. 11.**
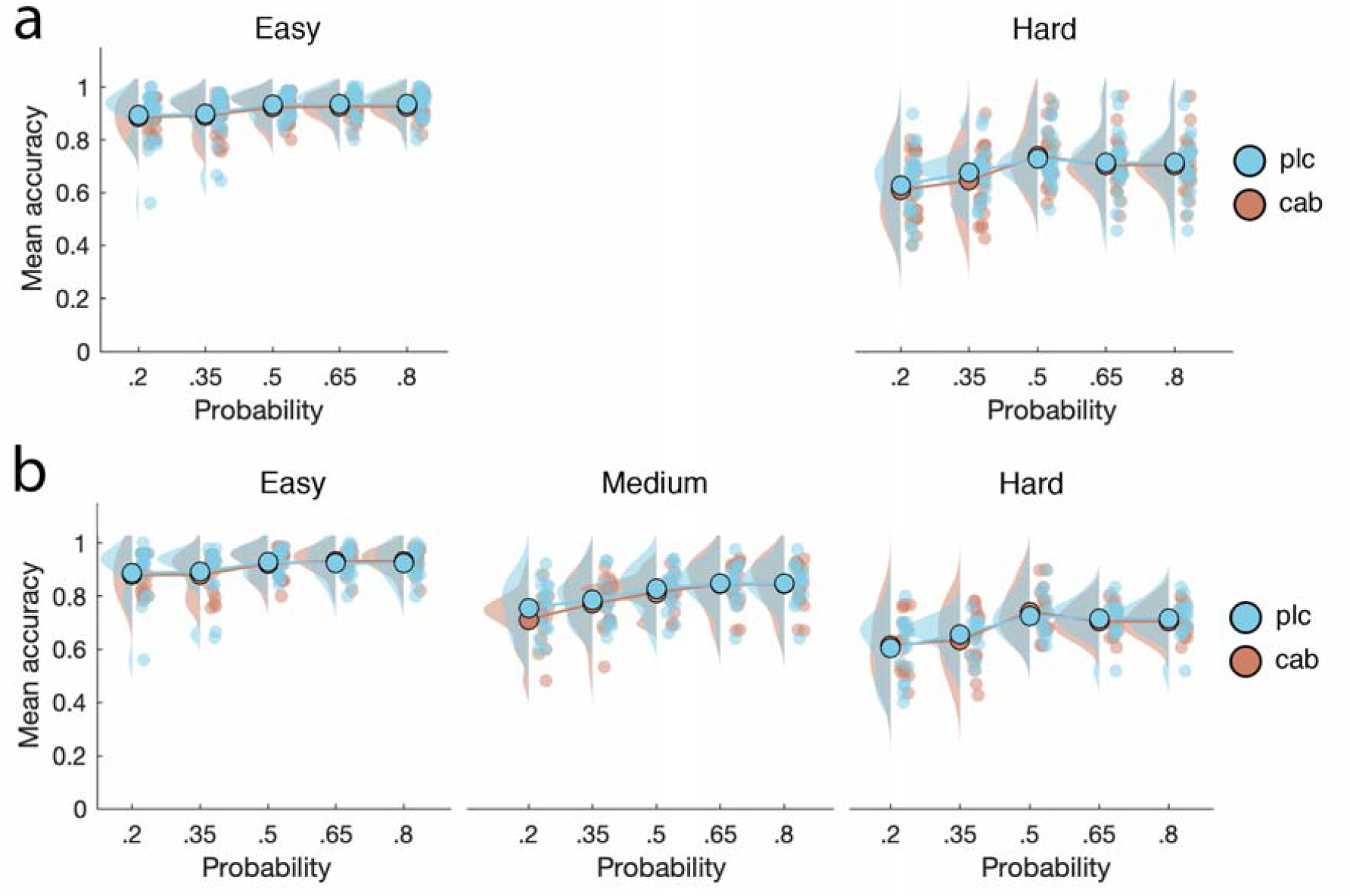
Cabergoline did not affect probabilistic discrimination performance. This figure shows mean accuracy for each drug condition and across five stimulus probabilities for all participants with two (**a**; N = 24) and three (**b**; N = 16) difficulty levels after titration. While accuracy was higher for easy trials and blocks in which the presented stimulus was more likely to occur, these effects were unaffected by cabergoline. Medium-difficulty trials take up position in between easy and hard trials in terms of mean accuracy when included

When we repeated these analyses for participants with all three difficulty levels for both stimuli after titration (N = 16; see Methods), the same pattern of results was obtained. Cabergoline did not affect discrimination accuracy (F_(1, 15)_ < 1, p = .47, BF_incl_ = .69), and we found no two-way interaction between Drug and Difficulty (F_(2, 30)_ < 1, p = .59, BF_incl_ = .05), Drug and Probability (F_(4, 60)_ < 1, p = .76, BF_incl_ = .02), or a three-way interaction (F_(8, 120)_ = 1.1, p = .4, BF_incl_ = .04; see Figure 11b). Accuracy was still lower on difficult (F_(2, 30)_ = 233.8, p < .001, η^2^ = .53, BF_incl_ > 100) and low-probability trials (F_(4, 60)_ = 21.1, p < .001, η^2^ = .08, BF_incl_ > 100). Controlling for screening sEBR and OSPAN did not change the acquired results. The interaction between drug Order and Drug was maintained (F_(1, 14)_ = 9.9, p = .007, BF_incl_ > 100), where overall accuracy was higher in the placebo condition (83±7%) when placebo was administered first compared to the cabergoline condition (80.6±6%). This difference was again absent when cabergoline was administered first (placebo: 82.7±3%, cabergoline: 83.1±3%).

Thus, we found no evidence in support of the conclusion that cabergoline affected any of the key findings in our perceptual tasks.

## Discussion

This double blind, placebo-controlled, cross-over study tested the hypothesis that striatal dopamine is involved in conscious perception by administering the dopamine D2 agonist cabergoline and placebo to healthy participants. To that end 1) we established an effect of cabergoline on participant’s physiological and subjective state, 2) we tested the effect of cabergoline on two often-used dopamine proxy measures (sEBR and OSPAN), and 3) we subjected participants to well-known and often-employed experimental paradigms targeting the neural correlates of consciousness. While we were able to establish an effect of cabergoline on participant’s physiological and subjective state, we did not find an effect of cabergoline on sEBR. Crucially, while we replicated key behavioral and ERP findings associated with the paradigms we employed (Del Cul et al., 2007; Slagter et al., 2017; Van Opstal et al., 2014), none of these findings were affected by cabergoline. Thus, we did not obtain evidence that the dopamine D2 agonist cabergoline affected conscious perception, which was also supported by Bayesian statistics.

Just as with positive results, an observed null result could indicate a true effect (i.e., no role for striatal dopamine in conscious perception), or it could be due to uncontrolled or unknown factors. Based on the convergent evidence discussed in the introduction implicating the striatum and its irrigation by dopamine in conscious perception (e.g., Bisenius et al., 2015; Slagter et al., 2017; Van Opstal et al., 2014), we predicted that cabergoline would affect performance on the backward masking and attentional blink tasks. Yet, our Bayesian results provided evidence against an effect of 1.5mg cabergoline (all BF_incl_ < .7; except for the interaction between Drug and drug Order for discrimination accuracy in the probabilistic discrimination task). This absence of an effect is further strengthened by the fact that other studies using different cognitive tasks have previously reported significant effects of cabergoline on task performance using a similar or an even lower dose of orally administered cabergoline (Broadway et al., 2018; Cavanagh et al., 2014; Fallon et al., 2017; Frank & O’Reilly, 2006; Norbury et al., 2013; Nandam et al., 2013; Yousif et al., 2016). These observations together argue for an interpretation in terms of a true null result. Nonetheless, there are a number of practical limitations that pertain to the present study.

For one, blinding was unsuccessful as 85% of participants guessed when they received cabergoline at the end of the experiment. While all participants with adverse reactions were excluded from any analyses, we cannot exclude the possibility that participants’ ability to tell when they received cabergoline influenced the results. It is possible this ability stems from the higher dosage (1.5 mg) we employed compared to previous studies administering cabergoline (1-1.25 mg; Broadway et al., 2018; Cavanagh et al., 2014; Frank & O’Reilly, 2006; Norbury et al., 2013; Nandam et al., 2013; Yousif et al., 2016). While a lower dosage may benefit blinding, it may also further reduce the chance of finding an effect of cabergoline. Another option for future research would be to administer an antiemetic such as domperidone in combination with cabergoline, in order to mitigate and mask physical side effects (Fallon et al., 2017; Norbury et al., 2013).

Second, while we titrated performance on the backward masking and probabilistic discrimination tasks, we did not do so for the attentional blink task, which may have resulted in reduced sensitivity of this task to our dopamine manipulation. We chose this specific version of the task in order to stay as close as possible to a recent study in which we found a relationship between activity in the ventral striatum and the attentional blink (Slagter et al., 2017). While we cannot rule out that a titration procedure may have resulted in a task more sensitive to our manipulation, we believe this is unlikely given that our participants showed a robust attentional blink, indicating that our task was suitable for the effect we intended to manipulate. In addition, since cabergoline did not affect performance on the two tasks we did titrate, we believe it unlikely that this would have been different in the case of the attentional blink task.

Third, a limitation of our study pertains to the use of alleged indirect measures of dopamine activity: sEBR (Jongkees & Colzato, 2016) and the OSPAN task (Cools, Gibbs, Miyakawa, Jagust, & D’Esposito, 2008). The functioning of dopamine relies on a relative equilibrium at the system level (i.e. frontal versus basal ganglia; Wiecki & Frank, 2013; Wise, Murray, & Gerfen, 1996), within the basal ganglia itself (i.e., direct versus indirect pathways; Redgrave et al., 2010), and at the level of synapses (i.e., pre-versus postsynaptic effects; Frank & O’Reilly, 2006; Usiello et al., 2000). The equilibrium at the system- and pathway-level is in turn implicated in the differing effects at the receptor level (i.e., D1 versus D2 receptors; Shen, Flajolet, Greengard, & Surmeier, 2008). Due to the complexity and the multitude of interactions between different systems it is especially important to be able to report on the initial state of the dopamine-system.

Unfortunately, sEBR was unaffected by cabergoline. While this finding goes against studies with humans showing effects of dopaminergic drugs on sEBR (Blin, Masson, Azulay, Fondarai, & Serratrice, 1990; Karson, 1983; Strakowski & Sax, 1998), this finding fits with several pharmacological studies that reported no effect (Depue, Luciana, Arbisi, Collins, & Leon, 1994; Ebert, 1996; Mohr, Sándor, Landis, Fathi, & Brugger, 2005; van der Post, de Waal, de Kam, Cohen, & van Gerven, 2004).

Moreover, differences in OSPAN and sEBR were unpredictive for the effect of cabergoline on performance in our experimental paradigms. These findings are surprising in light of previous work showing how effects of cabergoline on cognitive function depend on individual baseline sEBR (Cavanagh et al., 2014) and OSPAN score (Broadway et al., 2018). However, it should be emphasized that evidence for a relationship between dopamine-proxy measures and dopamine levels is often correlational, based on studies with small sample sizes (N < 50; Cools et al., 2008), and that results are mixed (Dang et al., 2017; Sescousse et al., 2018). It is debatable whether such small-sample studies have adequate power to provide evidence for this relationship (Cremers, Wager, & Yarkoni, 2017; Rousselet & Pernet, 2012). Similarly, the correlational findings we report should also be interpreted cautiously. While measures such as OSPAN and sEBR are easy to administer, the correlational evidence – as well as inconsistent findings regarding these measures (Dang et al., 2017; Jongkees & Colzato, 2016; Sescousse et al., 2018) – put pressure on their validity as an index of baseline dopamine levels.

Despite these practical limitations, the advantage of our study is that our results are univocal: the administration of cabergoline had no effect on any of our indices of conscious perception obtained using often-employed experimental paradigms in the study of consciousness. Of importance, we did consistently replicate all key behavioral and event-related potential (ERP) findings associated with each of these tasks.

Theoretically, perhaps our null-findings are best explained by an appeal to predictive processing approaches to brain function (Hohwy, 2012; 2013; de Lange, Heilbron, & Kok, 2018). From the perspective of these theories, the brain is viewed as a prediction machine which is continuously revising its predictions about the causes of sensory data. On this view, top-down signals in perceptual hierarchies carry perceptual predictions, while bottom-up signals convey perceptual prediction errors. Perception becomes a process of continual minimization of prediction errors across hierarchical levels, instantiating a process of approximate Bayesian inference on the causes of sensory signals. In this process, dopamine is thought to regulate the relative precision of top-down predictions and bottom-up prediction errors, where precision is understood as inverse variance (Friston et al., 2012): a higher (expected) variance of sensory signals leads to a smaller influence on updating perceptual predictions. “If true, this means that modulators of synaptic gain (like dopamine) do not report perceptual content but the context in which percepts are formed. In other words, dopamine reports the precision or salience of sensorimotor constructs (representations) encoded by the activity of the synapses they modulate.” (Friston et al., 2012, p. 2).

From this perspective, there may not have been enough uncertainty in our experimental paradigms for our dopamine manipulation to play a determining role. Indeed, previous studies employing cabergoline in the context of working memory found an effect under conditions where task-relevant targets were embedded in a larger perceptual field; namely, in the presence of distractors. In these studies, cabergoline exerted an effect on task performance in terms of target-detection accuracy and successful recall when a target stood in competition with other stimuli (Broadway et al., 2018; Cavanagh et al., 2014; Frank & O’Reilly, 2006). In the case of each of our tasks, targets were consistently surrounded by distracting information *temporally, but never spatially.* Perhaps the selective functionalities of the basal ganglia and dopaminergic firing actualize primarily when the system is under pressure to select sensorimotor constructs in the face of multidimensional uncertainty.

Our data cast doubt on a causal role for dopamine in visual perception, as shown by a significant lack of effect in several standard perceptual paradigms. Future studies, perhaps with more direct measures of dopaminergic activity, and more naturalistic paradigms including more opportunities for selection and/or uncertainty, may yet reveal a more specific influence of this neurotransmitter on how we perceptually encounter the world.

